# APRANK: computational prioritization of antigenic proteins and peptides from complete pathogen proteomes

**DOI:** 10.1101/2021.04.27.441630

**Authors:** Alejandro D. Ricci, Mauricio Brunner, Diego Ramoa, Santiago J. Carmona, Morten Nielsen, Fernán Agüero

## Abstract

Availability of highly parallelized immunoassays has renewed interest in the discovery of serology biomarkers for infectious diseases. Protein and peptide microarrays now provide a rapid, high-throughput platform for immunological testing and validation of potential antigens and B-cell epitopes. However, there is still a need for tools to prioritize and select relevant probes when designing these arrays. In this work we describe a computational method called APRANK (Antigenic Protein and Peptide Ranker) which integrates multiple molecular features to prioritize antigenic targets in a given pathogen proteome. These features include subcellular localization, presence of repetitive motifs, natively disordered regions, secondary structure, transmembrane spans and predicted interaction with the immune system. We applied this method to prioritize potentially antigenic proteins and peptides in a number of bacteria and protozoa causing human diseases: *Borrelia burgdorferi* (Lyme disease), *Brucella melitensis* (Brucellosis), *Coxiella burnetii* (Q fever), *Escherichia coli* (Gastroenteritis), *Francisella tularensis* (Tularemia), *Leishmania braziliensis* (Leishmaniasis), *Leptospira interrogans* (Leptospirosis), *Mycobacterium leprae* (Leprae), *Mycobacterium tuberculosis* (Tuberculosis), *Plasmodium falciparum* (Malaria), *Porphyromonas gingivalis* (Periodontal disease), *Staphylococcus aureus* (Bacteremia), *Streptococcus pyogenes* (Group A Streptococcal infections), *Toxoplasma gondii* (Toxoplasmosis) and *Trypanosoma cruzi* (Chagas Disease). We have tested this integrative method using non-parametric ROC-curves and made an unbiased validation using an independent data set. We found that APRANK is successful in predicting antigenicity for all pathogen species tested, facilitating the production of antigen-enriched protein subsets. We make APRANK available to facilitate the identification of novel diagnostic antigens in infectious diseases.

## 1 INTRODUCTION

Infectious diseases are one of the first causes of death worldwide, disproportionately affecting poor and young people in developing countries. Several epidemiological and medical strategies exist to deal with these diseases, most of which rely on robust and accurate diagnostic tests. These tests are used to demonstrate infection (presence of the pathogen), to follow up treatments and to monitor the evolution or cure of the disease or the success of field interventions (Peeling and Nwaka, 2011).

One of the preferred methods to diagnose infections relies on the detection of pathogen-specific antibodies in the fluids of infected patients (most often serum obtained from blood) (Washington, 1996; Vainionpää and Leinikki, 2008). However, knowledge of B-cell antigens and epitopes is scarce for many species. For this reason, there is a big interest in developing reliable methods able to improve the fast and sensitive identification of potential specific antigens.

With the advent of peptide microarray platforms it is now possible to perform high-throughput serological screening of short peptides, which allows for faster discovery of linear antigenic determinants with good potential for diagnostic applications (Pellois et al., 2002). Taking advantage of complete genome sequences from pathogens, it is theoretically possible to scan every encoded protein with short peptides against sera from infected hosts. However, while this is straightforwardly achieved for viral pathogens and small bacteria, it gets more difficult when dealing with larger bacteria or eukaryotic parasites, since they can reach thousands of proteins with millions of peptides, exceeding the average capacity of standard protein or peptide microarrays (Sutandy et al., 2013). Besides, it is now becoming common to fit in the arrays additional sequence variants obtained from the pathogen population (from diverse strains and clinical isolates). One example are serological strain typing strategies (Balouz et al., 2021), which would stress the capacity of these platforms.

Recently, ultrahigh-density peptide microarrays had been used successfully to map linear epitopes, having an upper theoretical limit of ~ 2-3 million unique peptides per array (Buus et al., 2012). While these ultrahigh-density peptide microarrays do enable a lot of possibilities, they do not yet have the capacity to analyze whole proteomes of larger pathogens without some preprocessing. It is also worth noting that they are not widely available as lower density arrays and they require substantial processing and downstream work to deal with large proteomes (Carmona et al., 2015; Durante et al., 2017; Mucci et al., 2017).

There are several ways to deal with the problem of not having enough space when accommodating large proteomes in a peptide array, each with their own advantages and disadvantages. The most common are: decreasing the overlap between peptides, dividing the proteome among different microarray slides, and using computational methods to prioritize antigens. In this paper we will focus on the latter. We and others have previously shown that a number of protein features can be used to validate and prioritize candidate antigens and epitopes for human pathogens (Carmona et al., 2012, 2015; Liu et al., 2018; Liang and Felgner, 2012). Similar approaches have also been developed into a number of reverse vaccinology programs for bacteria (reviewed recently in Dalsass et al. (2019)).

In a previous work, we developed a method that integrates information from a number of calculated molecular and structural features to compute an antigenicity score for proteins and peptides in *Trypanosoma cruzi* (Carmona et al., 2012, 2015). In this paper, we use machine learning techniques to extend and generalize this concept so that it can be applied to other pathogens. We call this method APRANK (Antigenic Protein and Peptide Ranker) and show how it can be used as a strategy to predict and prioritize diagnostic antigens for several human pathogens.

## 2 MATERIALS AND METHODS

### 2.1 Bioinformatic analysis

FASTA files containing proteins of the species used to train APRANK (see Table 1) were downloaded from publicly available database resources (from complete proteomes). To comply with requirements of downstream predictors, unusual amino acid characters were replaced by the character ‘X’ and a few proteins with more than 9,999 amino acids were truncated to that size. To obtain information at peptide level, proteins were split into peptides of 15 residues with an overlap of 14 residues between them (meaning an offset of 1 residue between peptides).

**Table 1.** List of pathogen species used in this paper.

| Pathogen Species | Disease | Group | Phylogenetic Lineage |
| --- | --- | --- | --- |
| *Borrelia burgdorferi* | Lyme disease | Gram Negative Bacteria | Spirochaetes |
| *Brucella melitensis* | Brucellosis |  | Alpha-proteobacteria |
| *Coxiella burnetii* | Q fever |  | Gamma-proteobacteria |
| *Escherichia coli* | Gastroenteritis |  | Enterobacteria |
| *Francisella tularensis* | Tularemia |  | Gamma-proteobacteria |
| *Leptospira interrogans* | Leptospirosis |  | Spirochaetes |
| *Porphyromonas gingivalis* | Periodontal disease |  | Sphingobacteria |
| *Mycobacterium leprae* | Leprosy | Gram Positive Bacteria | Actinobacteria |
| *Mycobacterium tuberculosis* | Tuberculosis |  | Actinobacteria |
| *Staphylococcus aureus* | Bacteremia |  | Firmicutes |
| *Streptococcus pyogenes* | GAS infections |  | Firmicutes |
| *Leishmania braziliensis* | Leishmaniasis | Eukaryotic Protozoa | Kinetoplastida |
| *Plasmodium falciparum* | Malaria |  | Apicomplexa |
| *Toxoplasma gondii* | Toxoplasmosis |  | Apicomplexa |
| *Trypanosoma cruzi* | Chagas Disease |  | Kinetoplastida |

Validated FASTA files were analyzed with BepiPred (Larsen et al., 2006), EMBOSS pepstats, Iupred (Dosztányi, 2018), NetMHCIIpan (Nielsen et al., 2010), NetOglyc (Julenius et al., 2005), NetSurfp (Klausen et al., 2019), Paircoil2 (McDonnell et al., 2006), PredGPI (Pierleoni et al., 2008), SignalP (Petersen et al., 2011), TMHMM (Krogh et al., 2001), Xstream (Newman and Cooper, 2007) and two custom perl scripts that analyzed similarity of short peptides against the human genome (NCBI BioProject PRJNA178030). The reasoning of choosing each predictor, what they predict and which version was used can be found in Table 2. The full console call for each predictor can be seen in Supplementary Table S2. NetMHCIIpan was run multiple times for different human alleles (DRB1*0101, DRB3*0101, DRB4*0101 and DRB5*0101). The only predictor that needed an extra preprocessing step was PredGPI, which required removing sequences shorter than 41 amino acids and those with an ‘X’ in their sequence. For all purposes, these filtered sequences were assumed to not have a GPI anchor signal. The versions of Linux, R, Perl, packages and modules used to create the computational method are listed in Supplementary Table S1.

**Table 2.**
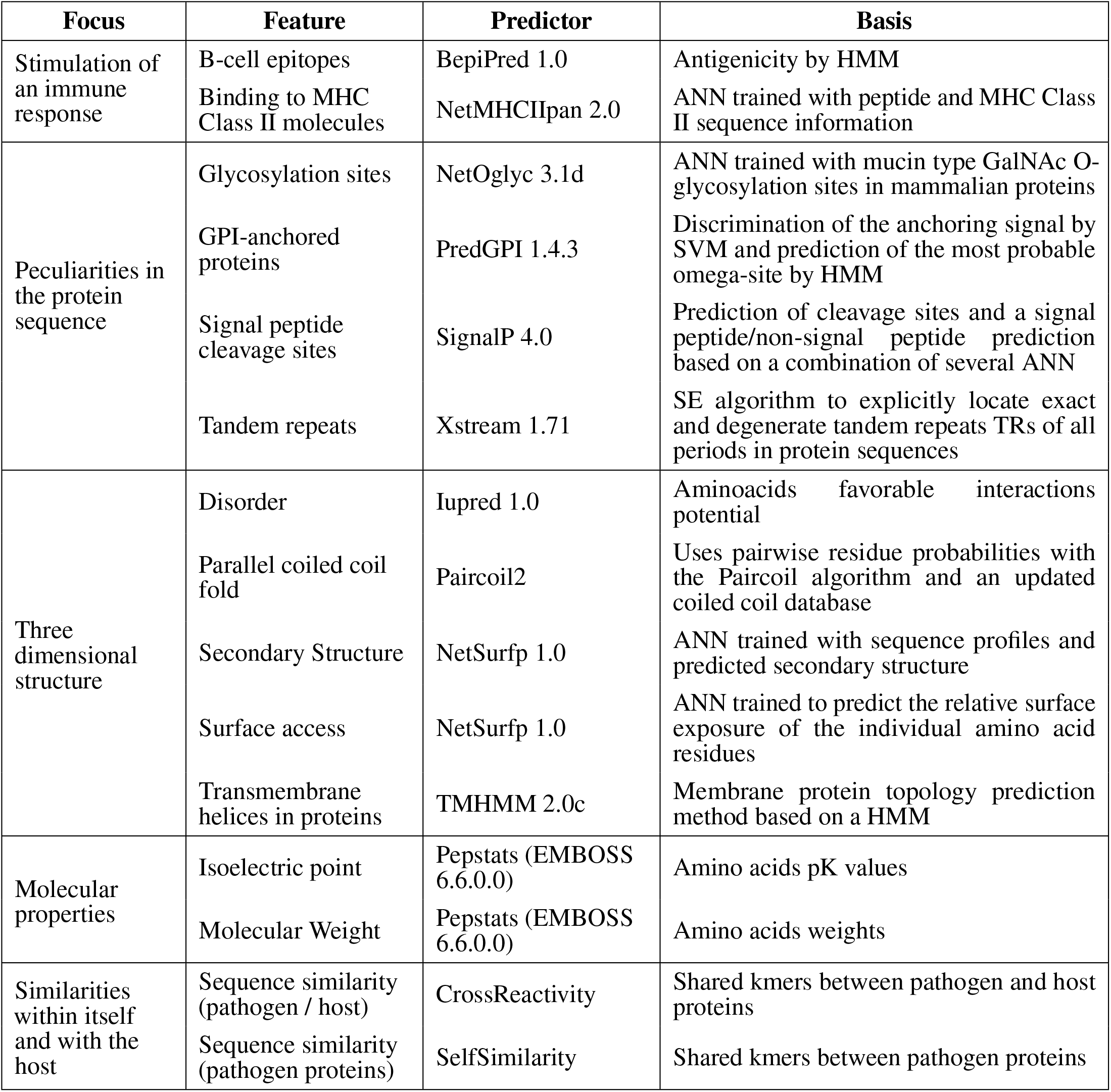
Predictors used to analyze different features of proteins and peptides. CrossReactivity and SelfSimilarity are custom Perl scripts. Acronyms used: ANN (Artificial Neural Network), HMM (Hidden Markov Model), SE (Seed Extension), SVM (Support Vector Machine).

### 2.2 Compiling a dataset of curated antigens

To obtain antigenic proteins and peptides, we extracted information from the immune epitope database (IEDB), as well as information from several papers, most of which relied on data from protein or peptide microarrays combined with sera of infected patients to find new antigens (Carmona et al., 2012; Vita et al., 2019; Martini et al., 2020; Xu et al., 2008; Barbour et al., 2008; Richer et al., 2015; Lawrenz et al., 1999; Eyles et al., 2007; Lu et al., 2007; Kilmury and Twine, 2010; Beare et al., 2008; Wang et al., 2013; Xiong et al., 2012; Vigil et al., 2011; Chen et al., 2009; Liang et al., 2010; Lessa-Aquino et al., 2013).

Because different protein identifiers are used across papers, we used either the Uniprot ID mapping tool, the blastp suite of BLAST or a manual mapping to find the corresponding ID or IDs that a given antigen had in our proteomes. The exhaustive list of all antigenic proteins and peptides used, their source and their mapping methods can be found in the Supplementary Data accompanying this article.

For the antigenic peptides, though, mapping the original protein ID to our pathogen proteomes was not enough; we also had to assign the antigenicity to the corresponding peptide or location within each antigenic protein, which meant dealing with the fact that the curated antigenic sequences varied in size. To do this, we developed our own mapping method that we called ‘kmer expansion’. This method marked as antigenic any peptide that shared a kmer of at least 8 amino acids with a curated antigenic sequence for the same protein. The amount of total and antigenic peptides, before and after the ‘kmer expansion’, are listed in Table 3.

**Table 3.** Amount of antigenic proteins and peptides for each species. This table shows the amount of antigenic proteins and sequences extracted from bibliography (to the left of the arrow) and the final amount after processing (to the right of the arrow). For proteins, BLAST was used to also tag as antigenic other proteins of the same species that were similar to the antigenic ones. For peptides, a custom mapping method named ‘kmer expansion’ was used to tag peptides as antigenic based on the antigenic sequences in bibliography (see Methods). We did not have information at peptide level for three of the species.

| Species | Group | Proteins |  | Peptides |  |
| --- | --- | --- | --- | --- | --- |
|  |  | Total | Antigenic | Total | Antigenic |
| *B. burgdorferi* | Gram - | 1,390 | 137 → 152 | 386,683 | 117 → 863 |
| *B. melitensis* | Gram - | 3,178 | 13 → 13 | - | - |
| *C. burnetii* | Gram - | 1,853 | 102 → 104 | - | - |
| *E. coli* | Gram - | 4,778 | 7 → 7 | 1,428,744 | 9 → 158 |
| *F. tularensis* | Gram - | 1,556 | 27 → 27 | - | - |
| *L. interrogans* | Gram - | 3,683 | 10 → 10 | 1,113,309 | 19 → 342 |
| *P. gingivalis* | Gram - | 1,881 | 10 → 11 | 626,536 | 165 → 1,181 |
| *M. leprae* | Gram + | 1,605 | 7 → 8 | 515,942 | 76 → 633 |
| *M. tuberculosis* | Gram + | 3,940 | 81 → 89 | 1,268,272 | 416 → 4,369 |
| *S. aureus* | Gram + | 2,607 | 16 → 16 | 758,970 | 55 → 575 |
| *S. pyogenes* | Gram + | 1,690 | 13 → 13 | 491,619 | 263 → 985 |
| *L. braziliensis* | Eukaryote | 8,084 | 8 → 12 | 4,964,396 | 14 → 182 |
| *P. falciparum* | Eukaryote | 5,337 | 106 → 131 | 4,009,580 | 562 → 9,120 |
| *T. gondii* | Eukaryote | 8,322 | 15 → 16 | 6,535,220 | 94 → 457 |
| *T. cruzi* | Eukaryote | 21,170 | 242 → 2,480 | 10,408,841 | 4,025 → 7,317 |

In the case of *Onchocerca volvulus*, the method we used to derive antigenic proteins and peptides was based on experimental proteome-wide data on antibody-binding to short peptides (Lagatie et al., 2017). We followed the same rules used by these authors and assigned as antigenic all peptides they called ‘immunoreactive’. Because in this work we are using an offset of 1 between overlapped peptides (maximum overlap), we also considered as antigenic the neighboring peptides that shared at least a kmer of length 8 with any immunoreactive peptide.

### 2.3 Clustering by sequence similarity

It is common practice in the literature to report antigenicity for a single or a few reference proteins or accession numbers. This information is then passed on to databases such as IEDB (Vita et al., 2019; Martini et al., 2020). Nevertheless, when dealing with complete proteomes, there are usually other paralogs with high sequence similarity to those labeled as antigenic. Since they have similar sequences, these proteins would then have similar properties which would likely result in similar outputs when running the predictors. However, because only one of those proteins is labeled as antigenic, this would hinder the learning capabilities of any models trained or tested with these data.

To improve the learning process of APRANK, and to account for unlabeled proteins, we calculated sequence similarity for all proteins in the 15 analyzed proteomes using blastp from the NCBI BLAST suite (Camacho et al., 2009) (console call in Supplementary Table S2). We then wanted to filter the BLAST output keeping only the good matches, which meant selecting a similarity threshold. After analyzing different matches, we arrived at a sensible compromise: trying to be as strict as possible without losing much data. For this we kept matches with a percentage of identical amino acids (pident) of at least 0.75, an expected value (evalue) less than or equal to 1 × 10^−12^ and a match length of at least half of the length of the shortest protein in the match.

Using these matches, we created a distance matrix where *distance* = 1 – *pident* and applied a single-linkage hierarchical clustering method. We then cut this tree using a cutoff of 0.25 (1 – *pidentThreshold*), resulting in a set of clusters of similar proteins.

For the species-specific models, proteins in a given cluster were kept together in the training process, meaning they would all be either in the training set or in the test set.

For the generic models, any protein in the training set which belonged to a cluster with at least one other antigenic protein was also tagged as antigenic, even across species (obviously excluding the species being tested). As for the test set in the generic models, this would also occur, but only inside that same species. The amount of total and antigenic proteins, before and after using BLAST to find similar proteins inside each species, can be see in Table 3.

### 2.4 Data normalization

Each predictor used by APRANK varied on how they returned their values. Not only they had different value ranges, but while some of them returned their values per protein, others did so per peptide, kmer, or amino acid. For this reason, we needed to parse and normalize all outputs before feeding their data into our model.

Values returned by each predictor were normalized to fit a numeric range between 0 and 1. Different methods were used to normalize the data depending on each predictor, ranging from linear or sigmoid normalizations to a simple binary indicator of presence or absence of a given feature (such as signal peptide). The detailed steps for the normalization at protein and peptide level for each predictor are described in Supplementary Table S3 and the formulas used for these operations can be found in Supplementary Article S1. The methods used to normalize the output for each predictor were the result of analyzing the distribution and spread of these outputs across all of our species for each predictor individually, coupled with biological knowledge of what each predictor was analyzing.

### 2.5 Fitting the species-specific models

Species-specific models were created to test the method and compare between balanced and unbalanced training sets. In this case a separate model was created for each species, using only train/test data from that organism alone. A schematic flowchart showing the logic of this procedure is shown in Figure 1. To fit each protein species-specific model, clusters for that species were divided in training and test sets in a 1:1 ratio due to the low number of recorded antigens for some species. For this same reason, the training set was balanced with ROSE (Lunardon et al., 2014), generating an artificial training set with a similar number of antigenic and non-antigenic artificial proteins. This process, as well as all other described below, was repeated 50 times by re-sampling the clusters in the training and test sets.

**Figure 1.**
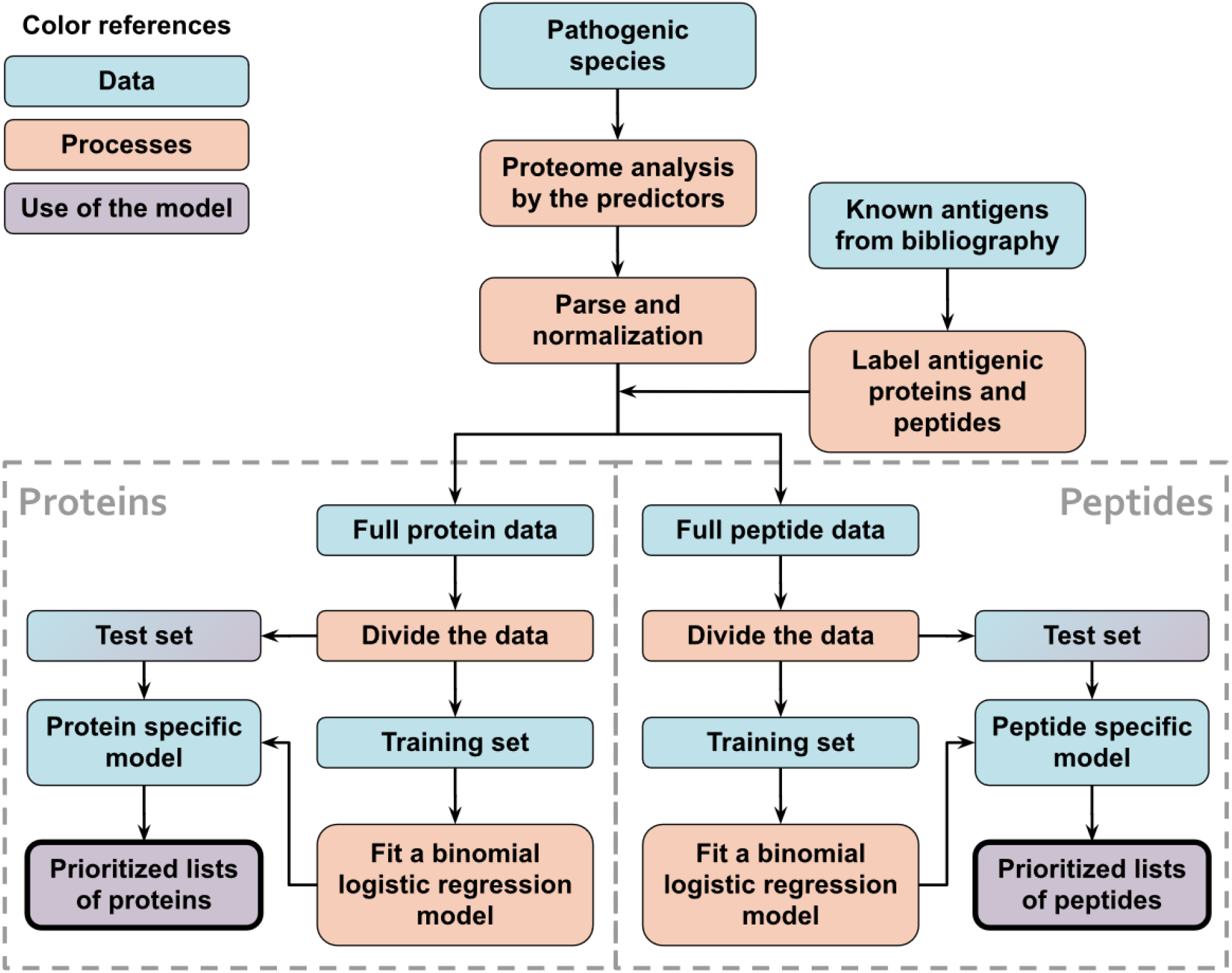
Schematic flowchart used to obtain APRANK’s species-specific models. With the aim of testing and tuning our method, training and prioritization was performed for both proteins and peptides using data from a single proteome of interest. This process was repeated for all of our 15 species.

A binomial logistic regression model was fitted for both the balanced and the unbalanced training sets using the generalized linear models in R (function `glm`). We chose this model for two reasons: because it allowed us to see a direct relationship between the models and our predictors via the coefficients of the model, and because it was not as affected as other more complex models by the existence of false negatives (which we knew existed because they were the novel antigens we wanted to find). Once the balanced and the unbalanced protein models were trained, we used them to predict the scores for the test set. The performance for each model, measured by the area under the ROC curve (AUC), was then calculated using the R package pROC (Robin et al., 2011). Additionally, two pseudo random set of scores were created by shuffling the scores achieved by both models. These random protein models were used to test if the performance of our models differed significantly from a random prediction.

For the peptide species-specific models, we divided the peptides into training and test sets by simply following the division of the proteins clusters, meaning that if a protein was in the training set for the protein model, its peptides would be in the training set for the peptide model for that iteration. The models were fitted and random scores calculated in a similar manner to the protein models. However, when we attempted to calculate the performance of the peptide models, our test set was too large to calculate performance based on AUC values in a reasonable time. We decided then to sample a subset of 50,000 peptides from the test set in a pseudo-random manner, making sure that the positive peptides were found in the subset and that the fraction of positive vs indeterminate/negative antigens was similar to the one in the test set (but never below 1% unless we ran out of antigens). All AUC values for the different peptide models were calculated using the same subset, and this process was repeated 5 times in each iteration, changing the subset each time.

Once all iterations were finished, we compared the AUCs obtained by the balanced and unbalanced models using a Student’s t-test. Another set of t-tests were used to analyze the difference between each of those models and their relative random model. If the model had a significantly higher AUC than the corresponding random model, we considered the model achieved a successful prediction (p < 0.05).

### 2.6 Creating the generic models

The generic (pan-species) models are the actual models used by APRANK. The objective of these models is to predict antigenic proteins and peptides for new species (which APRANK have never seen before). In a broad sense, they have to understand what makes a protein or a peptide antigenic. We achieved this by training the models with a large set of antigenic proteins and peptides from 15 different species, including gram-positive bacteria, gram-negative bacteria and eukaryotic protozoans.

To create the protein generic models, we used ROSE (Lunardon et al., 2014) to create a balanced training set of 3,000 proteins for each species and then merged all those balanced training sets together. With these data, a linear model was created following the same steps as for the species-specific models. Next, these models were used to predict the scores for the species being analyzed and the performance of the prediction was calculated the same way as for the species-specific protein models. A schematic visualization of this procedure is shown in Figure 2.

**Figure 2.**
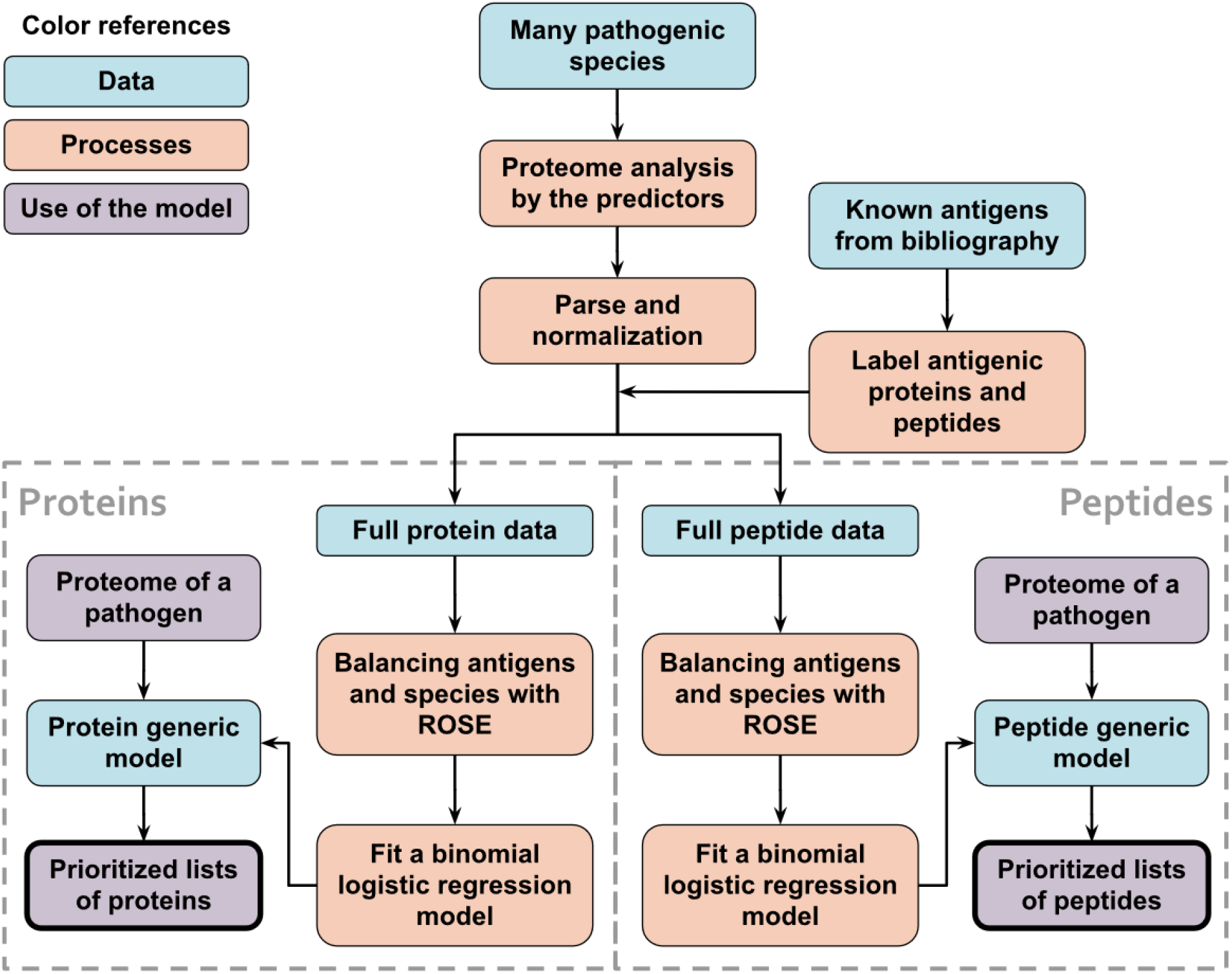
Schematic flowchart used to obtain APRANK’s generic models. With the aim of creating a model that could make predictions for a wide range of species, training and prioritization was performed for both proteins and peptides using combined data from all of our 15 species. When testing these models, leave-one-out models were used, where 14 species were used to train the model and the 15th species to test it. This process was repeated for all of our 15 species.

We created the peptide generic models in a similar manner, with balanced training sets from each of the species that contained 100,000 peptides each. In addition to the regular score calculated by using the model to predict the antigenicity of the test data, we also calculated a combined score, which was simply the mean of the protein and peptide scores for that peptide. The performance of the peptide generic models was calculated the same way as for the species-specific peptide models.

When testing these generic models, we created temporary leave-one-out generic models, where we used 14 of the species to generate the model, and then tested the model in the 15th species. We then generated the final protein and peptide generic models using all 15 species and tested them by predicting antigenicity in *Onchocerca volvulus*, a novel species with experimental proteome-wide data (Lagatie et al., 2017).

### 2.7 Comparative performance

To discard the possibility that our model was simply detecting sequence similarity, we created a ‘BLAST model’, where we assigned to each protein a score based solely on how similar they were to a known antigenic protein from another organism. The score used was – *log*_10_(*evalue*) and then performance was calculated for each species.

We also wanted to make sure our model was combining information from several predictors. To rule out that performance was mainly driven by one predictor, we compared our prediction capabilities against the individual predictor with best AUC, which was BepiPred 1.0. To do this, the BepiPred score for each protein and peptide was obtained from the individual amino acid scores following the same steps we used in APRANK as detailed in Supplementary Table S3, but without normalizing it. The AUCs for the BepiPred peptide scores were calculated the same way as for the peptide species-specific models.

### 2.8 Availability

The code for running or modifying APRANK is available at GitHub (Ricci and Agüero, 2021), released under a BSD 2-Clause ‘Simplified License’, which is a permissive free software license. The repository also holds documentation on how to configure, and install dependencies (users are responsible for obtaining the corresponding licenses or permissions for some required predictors); as well as the trained generic models for proteins and peptides (in R files of type .*rda* containing compressed data structures).

## 3 RESULTS

Our aim in this work was to develop a computational method and associated pipeline capable of prioritizing candidate antigenic proteins and antigenic determinants (epitopes) from complete pathogen proteomes for downstream experimental evaluation. We have previously shown for *Trypanosoma cruzi* (Chagas Disease) that different criteria can be integrated and exploited in a computational strategy to further guide the process of diagnostic peptide discovery (Carmona et al., 2012). Here we extend this work to other human pathogens and improve the way in which features are weighted, hence providing a tool for the prioritization of candidate linear B-cell epitopes for a wide range of pathogens.

### Species and Features

We selected human pathogens from a phylogenetically diverse set of taxa with experimentally validated antigen and/or epitope data to train and test our method. This included gram negative bacteria, gram positive bacteria and eukaryotic protozoans. The species and the diseases they cause are shown in Table 1.

We obtained the proteomes of these species (see Methods) and split each protein into peptides of 15 residues. Once this was done, we used information from the immune epitope database (IEDB) along with manually extracted information from several papers to tag each protein and peptide as antigenic or non-antigenic. The ‘non-antigenic’ tag in this paper should be understood in the sense of proteins with no prior information on their antigenicity. The amount of total and antigenic proteins and peptides can be see in Table 3.

To develop a tool that can help identify candidate antigenic proteins and peptides, we used several predictors that focused on different properties of the proteins (Table 2). On a broad sense, these predictors assess: the antigenicity and/or immunogenicity of proteins (Larsen et al., 2006; Nielsen et al., 2010); the structural and post-translational features that can be predicted from the protein sequence, some of which may suggest the protein enters the secretory route or is anchored at the membrane (Julenius et al., 2005; Pierleoni et al., 2008; Petersen et al., 2011); the presence of internal tandem repeats in proteins, which have been described to modulate immunogenicity of proteins (Newman and Cooper, 2007) together with other structural features such as the presence of intrinsically unstructured or exposed regions in proteins which may effect their presentation in the context of an immune response (Dosztányi et al., 2005; McDonnell et al., 2006; Petersen et al., 2009; Krogh et al., 2001).

We have also implemented in APRANK a number of custom Perl and R scripts that measure sequence similarity between each pathogen protein and the human host (CrossReactivity), or itself (SelfSimilarity). The idea behind these measurements was to obtain additional information on highly conserved sequences that may result in e.g. potential lack of immune response (tolerance) if the pathogen sequence is highly similar to a human protein; or cross-reactivity of antigens and epitopes in other proteins from the same pathogen (self-similarity). These predictors provide information on desirable and undesirable properties that then need to be weighted accordingly to achieve good performance at the task of antigen and epitope prediction.

### Testing APRANK and ROSE on species-specific models

Species-specific models were created to test the method and to compare between unbalanced training sets and training sets balanced using ROSE (see Methods). As the name implies, these models worked with only one species at a time, using a fraction of its proteins to predict antigenicity for the rest. After running the predictors for all proteins in the selected genome, we parsed and processed the different outputs and applied a normalization process to have them in a common scale.

We needed to divide our data into training and test sets. Often, training sets represent ~ 80% of the data; however, in our case some species had a low number of validated antigens (see Table 3), which meant that choosing a 80/20 training/test set split would result in test sets having only a few antigenic proteins. This kind of imbalance tends to compromise the training process, making the model to focus on the prevalent class (non-antigenic) and ignore the rare class (antigenic) (Menardi and Torelli, 2014). For this reason, when training a model using data from a single species, we chose to split the training and test set 50/50, re-sampling proteins and peptides multiple times (see Methods). To improve the training process, we also used ROSE to balance our training sets, which works by generating artificial balanced samples from the existing classes, according to a smoothed bootstrap approach (Lunardon et al., 2014). Furthermore, we used the similarity-based clustering of sequences to avoid placing highly similar sequences into both training and test sets.

We used these balanced training sets to fit a binomial logistic regression model, resulting in one model for proteins and one for peptides. These models, which we denominated *species-specific models*, were then used to predict the antigenicity of their respective test sets. The performance of APRANK was assessed by measuring the area under the ROC curve (AUC), using known antigens and epitopes in the protein and peptide test sets. This whole process was repeated 50 times, re-sampling which proteins were in the training set and which in the test set. A final APRANK AUC score for each species was calculated as the mean of all AUC scores for these iterations (see Figure 3). To assess the effect of balancing the data on our models using ROSE, we also assessed the performance of APRANK repeating the procedure described above using the unbalanced training sets instead, resulting in a set of AUC scores corresponding to species-specific models trained with unbalanced data.

**Figure 3.**
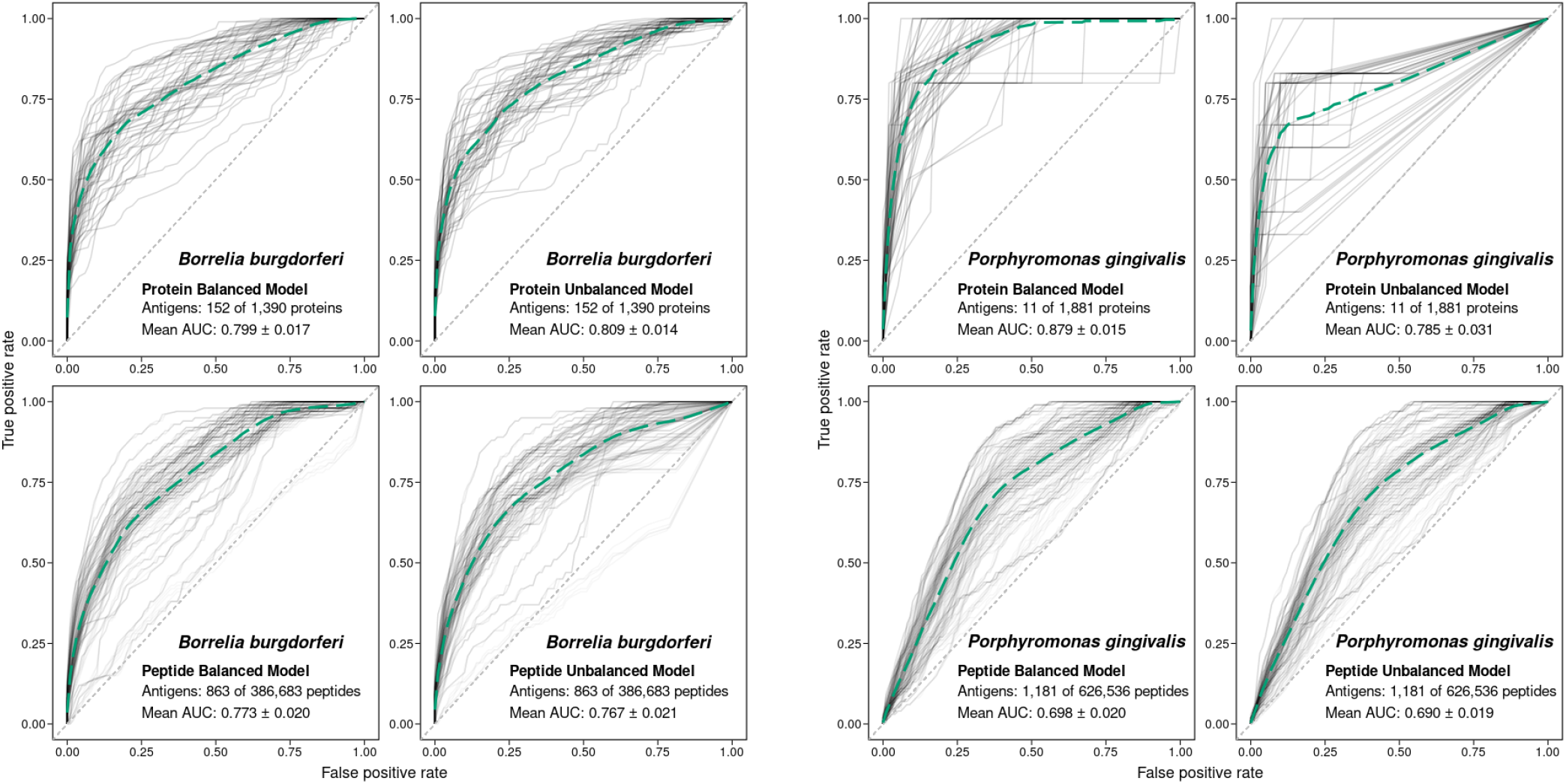
Performance of APRANK training using balanced or unbalanced data. Performance of APRANK’s species-specific models for *B. burgdorferi* and *P. gingivalis*. ROC curves for each iteration of training and testing are shown in light gray, and the average curves are shown in green (dashed lines).

These calculations were done for each of the 15 species, although for 3 of them there was no antigenicity information at the peptide level, and only protein models were calculated. The results are presented in Table 4. Our testing showed that APRANK was able to predict antigenicity for proteins and peptides in most cases, with good performance. The only species that did not have a successful prediction were *E. coli* for the protein model, and *M. tuberculosis* and *S. aureus* for the peptide model. In these cases, the final AUC corresponding to the species-specific model was not significantly different than a random prediction. As for the balancing of the data using ROSE, it seemed to have mostly positive or neutral effects in the predicting capabilities of our models, which meant we could safely use it in training our pan-species models.

**Table 4.** Prediction results for the specific models. The prediction was considered to be successful if it was significantly Better Than a Random set of scores (BTR). Each specific model was calculated 50 times using different, but overlapping, subsets of data as training and test sets. In bold we show the model with the significantly higher AUC when comparing training with unbalanced or balanced data (Student’s t-test, * < 0.05, ** < 0.01, *** < 0.001).

| Species | Proteins |  |  | Peptides |  |  |
| --- | --- | --- | --- | --- | --- | --- |
|  | BTR | Trained with unbalanced data | Trained with balanced data | BTR | Trained with unbalanced data | Trained with balanced data |
|  |  | Mean AUC | Mean AUC |  | Mean AUC | Mean AUC |
| B. burgdorferi | Yes | 0.809 ± 0.014 | 0.799 ± 0.017 | Yes | 0.767 ± 0.021 | 0.773 ± 0.020 |
| B. melitensis | Yes | 0.710 ± 0.037 | 0.700 ± 0.033 | - | - | - |
| C. burnetii | Yes | 0.611 ± 0.011 | 0.620 ± 0.010 | - | - | - |
| E. coli | **No** | 0.511 ± 0.034 | 0.515 ± 0.039 | Yes | 0.584 ± 0.056 | 0.633 ± 0.047 |
| F. tularensis | Yes | 0.783 ± 0.018 | **0.807 ± 0.014\*** | - | - | - |
| L. interrogans | Yes | 0.827 ± 0.033 | 0.867 ± 0.023 | Yes | 0.559 ± 0.015 | 0.565 ± 0.011 |
| P. gingivalis | Yes | 0.785 ± 0.031 | **0.879 ± 0.015\*\*\*** | Yes | 0.690 ± 0.019 | 0.698 ± 0.020 |
| M. leprae | Yes | 0.633 ± 0.018 | 0.652 ± 0.018 | Yes | 0.557 ± 0.029 | 0.585 ± 0.023 |
| M. tuberculosis | Yes | 0.635 ± 0.010 | 0.647 ± 0.011 | **No** | 0.508 ± 0.010 | 0.502 ± 0.010 |
| S. aureus | Yes | 0.765 ± 0.032 | 0.772 ± 0.023 | **No** | 0.438 ± 0.054 | 0.420 ± 0.057 |
| S. pyogenes | Yes | 0.884 ± 0.039 | **0.984 ± 0.003\*\*\*** | Yes | 0.832 ± 0.021 | 0.844 ± 0.019 |
| L. braziliensis | Yes | **0.719 ± 0.021\*\*** | 0.673 ± 0.020 | Yes | 0.778 ± 0.029 | **0.867 ± 0.025\*\*\*** |
| P. falciparum | Yes | 0.821 ± 0.009 | 0.826 ± 0.007 | Yes | 0.758 ± 0.016 | **0.779 ± 0.012\*** |
| T. gondii | Yes | 0.656 ± 0.032 | **0.744 ± 0.032\*\*\*** | Yes | **0.646 ± 0.035\*\*** | 0.584 ± 0.020 |
| T. cruzi | Yes | 0.803 ± 0.029 | **0.850 ± 0.022\*** | Yes | 0.838 ± 0.019 | 0.854 ± 0.016 |

### Development of APRANK as a pan-species ranker of antigens and epitopes

In the previous section we used protein and peptide data from a given pathogen species to train a model that successfully predicted antigenicity for that same organism; however, our end goal was to have a model that was able to predict antigenicity for any pathogen. To achieve this, we created models trained with all species, which we called *protein generic models* and *peptide generic models*.

For these models, we used ROSE (Lunardon et al., 2014) to generate similar sized partitions of balanced data for each of the species, and then we merged this data and fitted a binomial logistic regression model, using the same as described before. When using the models to predict the peptide antigenicity scores, we also analyzed the predicting capabilities of what we called the *combined score*, which was a combination of the protein and peptide scores for a given peptide.

To validate these models we performed a leave-one-out cross-validation method (LOOCV), hence creating 15 different protein generic models, each time leaving out one species (which was the one being used as test set). For the peptide generic models we followed a similar route, but we ended up with 12 models due to the lack of antigenicity information at peptide level for 3 of the 15 species.

The performance results for these models are presented in Table 5. The generic protein models were successful in predicting antigenicity for all species, and similar results were obtained also at the peptide level, achieving successful predictions even for *E. coli*, *M. tuberculosis* and *S. aureus*, which were the three species where the species-specific models performed poorly before. This observation suggests that performance is related to the amount and diversity of recorded antigens. As for the performance of these generic models, the observed AUC scores obtained similar values to the ones obtained in the species-specific models trained with balanced data, indicating that while these generic models did not have information about the species being tested, the data obtained from all the other 14 species was enough to learn the generic rules that made a protein antigenic.

This is also evident when comparing the coefficients obtained in the different protein models. In the case of individual (species-specific) models, coefficients were less robust across iterations when there were few positive cases, and more robust with larger validated training examples, as expected (see Supplementary Figures S1 and S2). For the pan-species models, we found the coefficients to be very robust across all 15 models, indicating that the different leave-one-out generic models reached a similar conclusion on what makes a protein ‘antigenic’ (see Supplementary Figure S3). This reinforces the idea that better performance is the result of more extensive training with diverse positive and negative examples.

**Table 5.** Prediction results for the leave-one-out generic models. The prediction was considered successful if it was significantly Better Than a Random set of scores (BTR). For peptides, we show both the performance of the model alone, and the performance obtained by combining the protein and peptide scores. In bold we show any difference greater than 5% between the peptide score and the combined score for a given species. LOO Model = Leave-One-Out Model.

| Species | Proteins |  | Peptides |  |  |  |
| --- | --- | --- | --- | --- | --- | --- |
|  | BTR | LOO model | BTR | LOO model | LOO model + protein scores | Combined score relative AUC gain |
| B. burgdorferi | Yes | 0.786 | Yes | 0.768 | 0.950 | **23.60%** |
| B. melitensis | Yes | 0.774 | - | - | - | - |
| C. burnetii | Yes | 0.620 | - | - | - | - |
| E. coli | Yes | 0.754 | Yes | 0.742 | 0.780 | **5.12%** |
| F. tularensis | Yes | 0.698 | - | - | - | - |
| L. interrogans | Yes | 0.947 | Yes | 0.679 | 0.948 | **39.57%** |
| P. gingivalis | Yes | 0.854 | Yes | 0.665 | 0.871 | **30.91%** |
| M. leprae | Yes | 0.758 | Yes | 0.692 | 0.731 | **5.68%** |
| M. tuberculosis | Yes | 0.702 | Yes | 0.586 | 0.711 | **21.17%** |
| S. aureus | Yes | 0.737 | Yes | 0.752 | 0.790 | **5.03%** |
| S. pyogenes | Yes | 0.983 | Yes | 0.838 | 0.970 | **15.81%** |
| L. braziliensis | Yes | 0.709 | Yes | 0.946 | 0.878 | **-7.20%** |
| P. falciparum | Yes | 0.807 | Yes | 0.748 | 0.835 | **11.66%** |
| T. gondii | Yes | 0.837 | Yes | 0.583 | 0.720 | **23.51%** |
| T. cruzi | Yes | 0.867 | Yes | 0.843 | 0.857 | 1.58% |

### Using APRANK to obtain antigen-enriched sets

Our generic models allowed us to rank proteins and peptides in a given species based on a model trained from other pathogens. Now, we wanted to use these scores to select a subset of proteins or peptides with an increased chance of being antigenic when compared to the whole proteome.

For this, we focused on *T. cruzi*, as this was the species with the largest number of recorded antigens within our collection. To obtain fair antigenicity scores for this protein we used the corresponding leave-one-out models created when testing the generic models. We analyzed the distribution of the normalized scores returned by these models, distinguishing between antigenic and non-antigenic proteins and peptides (see Figure 4). As was expected, the peak of the scores for the antigens is found to the right of the one for the non-antigens, indicating that the average score is higher for the antigenic proteins and peptides. Also, the amount of overlapping can be related to the corresponding AUC, where the higher the AUC, the less the overlapping.

**Figure 4.**
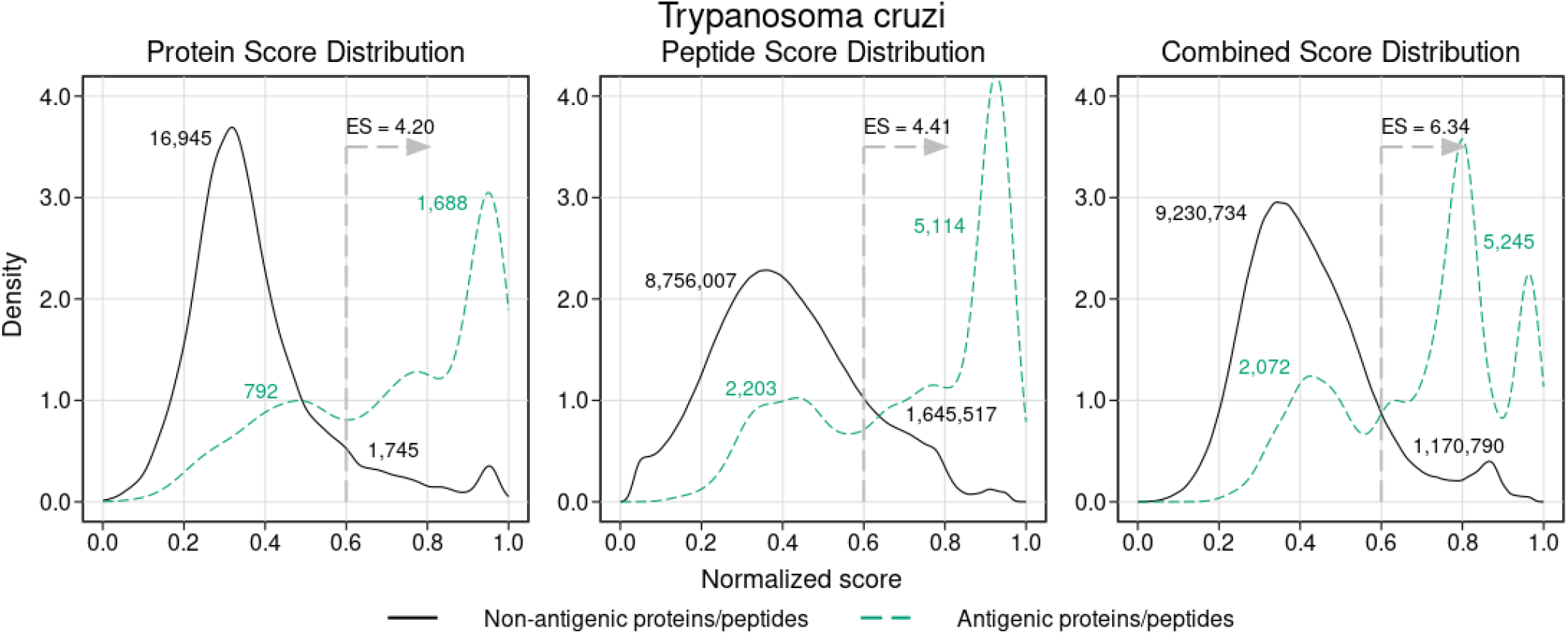
Density analysis for the antigenicity scores of *T. cruzi*. Plots were obtained by analyzing the proteome of *T. cruzi* with the leave-one-out generic models, and then distinguishing between antigens and non-antigens. The figure shows the enrichment score obtained by keeping only the proteins and peptides with a score greater than 0.6, as well as the amount of antigens and non-antigens that would be inside or outside that subset.

Once we had our score distributions, we used them to select an antigen-enriched subset of proteins and peptides. This could be done in one of two ways: either by setting a score threshold or by simply selecting a fixed number of proteins and peptides within the top scores. After analyzing the distribution of score values, we decided to use the first option and selected those proteins and peptides with a normalized score of at least 0.6. We next calculated what we called *enrichment score* (ES), which was the proportion of antigens in the selected subset relative to the proportion of antigens in the whole proteome (for example, ES = 2 meant you were twice as likely to find an antigen in the subset than in the whole proteome, or in a random subset). In Figure 4 we show the enrichment scores for the different normalized scores and the number of proteins and peptides that fall inside or outside those subsets. While the subsets were usually a small fraction of the whole proteome (close to 10% in most cases), this represents a 4 – 6 fold increase in the chances of finding antigens in those subsets.

As an example, suppose a microarray with a capacity of 200,000 unique peptides. Based on the current antigenic data we possess, a random sampling of the *T. cruzi* proteome would lead to the inclusion of ~ 140 antigenic peptides in that microarray. However, using APRANK to select the top 200,000 peptides with the highest normalized combined score, we would end up including almost 1,600 antigenic peptides in the array (an enrichment score of 11.35). This demonstrates the utility of tools like APRANK for selection of antigenic peptides for screening platforms.

### Assessing the validity of the computational method

Now that we had a working pan-species model, we next analyzed the contribution of each predictor to the overall predicting capabilities of APRANK. This was done to confirm that the performance achieved by APRANK came from combining information from different predictors, and not from just one or a few of them. For this, we calculated the predicting capabilities of each individual predictor using their output as score (data not shown). We found that the predictor with best solo predicting capabilities was BepiPred 1.0, so we compared its predictions against APRANK’s for both the protein and peptide generic models for each species, which can be seen in Table 6.

**Table 6.** Comparison between APRANK and the predictor with highest solo AUC (BepiPred 1.0). The relative AUC gain shows the increase or decrease of the AUC obtained by our method relative to the one obtained by BepiPred. Differences greater than 5% are shown **in bold**.

| Species | Proteins |  |  | Peptides |  |  |
| --- | --- | --- | --- | --- | --- | --- |
|  | BepiPred score AUC | APRANK score AUC | APRANK relative AUC gain | BepiPred score AUC | APRANK score AUC | APRANK relative AUC gain |
| B. burgdorferi | 0.729 | **0.786** | **7.94%** | 0.796 | 0.768 | -3.46% |
| B. melitensis | 0.710 | **0.774** | **8.93%** | - | - | - |
| C. burnetii | 0.558 | **0.620** | **11.13%** | - | - | - |
| E. coli | 0.587 | **0.754** | **28.39%** | 0.662 | **0.742** | **12.21%** |
| F. tularensis | 0.570 | **0.698** | **22.40%** | - | - | - |
| L. interrogans | 0.839 | **0.947** | **12.87%** | 0.676 | 0.679 | 0.42% |
| P. gingivalis | 0.852 | 0.854 | 0.25% | 0.674 | 0.665 | -1.36% |
| M. leprae | **0.868** | 0.758 | **-12.67%** | 0.689 | 0.692 | 0.51% |
| M. tuberculosis | 0.666 | **0.702** | **5.29%** | 0.561 | 0.586 | 4.58% |
| S. aureus | 0.723 | 0.737 | 1.86% | 0.767 | 0.752 | -1.93% |
| S. pyogenes | 0.970 | 0.983 | 1.33% | 0.8 | 0.838 | 4.73% |
| L. braziliensis | 0.549 | **0.709** | **29.00%** | 0.905 | 0.946 | 4.48% |
| P. falciparum | 0.793 | 0.807 | 1.84% | 0.642 | **0.748** | **16.42%** |
| T. gondii | 0.579 | **0.837** | **44.59%** | 0.584 | 0.583 | -0.21% |
| T. cruzi | 0.814 | **0.867** | **6.54%** | 0.819 | 0.843 | 3.03% |

We focused on those cases where the AUC changed at least 5% between BepiPred 1.0 and APRANK’s generic models. APRANK showed increased predicting capabilities for 11 out of the 15 analyzed proteomes at the level of complete proteins and/or peptides, while showing a decrease in performance only in *M. leprae* at protein level. These results provide validation support to the approach built into APRANK by combining information from many predictors.

As an additional test, we also assessed the performance of APRANK after removing BepiPred 1.0 predictions from our model. This can be seen in Supplementary Table S4. In this simulation we observed that even without BepiPred 1.0, our model reached similar predicting capabilities in most cases, hence suggesting that other predictors and features included in APRANK were able to replace BepiPred when training the model (this is further discussed in the Conclusions).

To ensure that our model was doing more than simply detecting sequence similarity, we also compared our performance against a ‘BLAST model’, meaning a model that was based solely on how similar a given protein was to a known antigenic protein. The comparison between the performance of this model and APRANK can be seen in Supplementary Table S5. As expected, APRANK achieved a larger AUC for most for the species; however we observed that for *M. leprae* and *L. braziliensis* the ‘BLAST model’ actually resulted in a better prediction. We believe this was due to them being species with a small amount of antigens and a high similarity to other of our selected species. To test this, we repeated this analysis for these two species, but now removing from the BLAST results (and so, from the ‘model’) the species that was the most similar to the one being analyzed. These new predictions indeed resulted in a considerable lower AUC, matching or falling behind APRANK.

### Applying our method on a novel species

As a final step, we wanted to test APRANK on a new species that was not included in our initial training and that had an extensive amount of information on the antigenicity of its proteins and peptides. For this, we searched for publications containing proteome-wide linear epitope screenings using high-density peptide microarrays and selected a recent dataset produced by scanning the complete *Onchocerca volvulus* proteome with more than 800,000 short peptides (mostly 15mers) (Lagatie et al., 2017). *Onchocerca volvulus* is a nematode and it is the causative agent of Onchocerciasis in humans (also called river blindness), a disease that is on the list of Neglected Tropical Diseases (NTDs) of the World Health Organization (Holmes, 2014).

To obtain a list of antigens in *O. volvulus*, we followed the same rules applied by the authors to find the peptides they called ‘immunoreactive’ (see Methods in Lagatie et al.), resulting in a set of almost 1,100 antigenic peptides. We tagged a protein as antigenic if it had at least one of these peptides; however, we also kept information on how many ‘immunoreactive’ peptides each protein had for later analysis. Once this was done, we also tagged as antigenic any neighboring peptide that shared at least 8 amino acids with one of these ‘immunoreactive’ peptides.

We next trained APRANK with all our 15 species and then used this model to predict the antigenicity scores for both the proteins and the peptides of *O. volvulus*. An AUC score was calculated for each prediction, comparing the score given by APRANK against the antigenic tag for each protein and peptide. We also calculated the enrichment scores for these scenarios using a score threshold of 0.6 in a similar way that we did for *T. cruzi*.

Our method was successful in predicting the antigenicity of proteins and peptides for *O. volvulus*, as shown in Table 7. We observed that if we were more strict when tagging a protein as antigenic, meaning requiring at least 2 or 3 ‘immunoreactive’ peptides before doing so, we obtained better performance. When considering as antigenic any protein with 1 ‘immunoreactive’ peptide we had an enrichment score of 2.28, whereas when we increased this requirement to 3 ‘immunoreactive’ peptides the enrichment score was 5.29 (see Table 7, Figure 5). Besides validating the performance of APRANK on a new pathogen, this suggests that either our method is better in predicting proteins with many antigenic regions, or that a single reactive peptide from a peptide array screening may provide only weak support for calling of antigens.

**Table 7.** Performance of APRANK on *Onchocerca volvulus*. Proteins and peptides were tagged as antigenic based on the number of Minimum Immunoreactive Peptides (#MIP). For proteins, we considered as antigenic those with at least #MIP immunoreactive peptides. For peptides, we considered as antigenic any immunoreactive peptide found inside proteins with at least #MIP immunoreactive peptides, and their neighboring peptides. The rule to define an ‘immunoreactive peptide’ was extracted from Lagatie et al. 2017 (see Methods). The enrichment score represents the proportion of antigens in the selected subset relative to the proportion of antigens in the whole proteome.

|  | Total | Score | #MIP | Antigenic | AUC | Antigens with score > 0.6 | Enrichment score for 0.6 |
| --- | --- | --- | --- | --- | --- | --- | --- |
| **Proteins** | 12,994 | Protein score | 1 | 886 | 0.677 | 150 | 2.28 |
|  |  |  | 2 | 177 | 0.713 | 38 | 2.89 |
|  |  |  | 3 | 28 | 0.828 | 11 | 5.29 |
| **Peptides** | 4,872,082 | Peptide score | 1 | 1,097 → 12,917 | 0.800 | 5,520 | 3.29 |
|  |  |  | 2 | 397 → 4,145 | 0.798 | 1,779 | 3.30 |
|  |  |  | 3 | 104 → 1,107 | 0.836 | 547 | 3.80 |
|  |  | Combined score | 1 | 1,097 → 12,917 | 0.750 | 3,035 | 3.05 |
|  |  |  | 2 | 397 → 4,145 | 0.774 | 1,189 | 3.73 |
|  |  |  | 3 | 104 → 1,107 | 0.871 | 478 | 5.61 |

**Figure 5.**
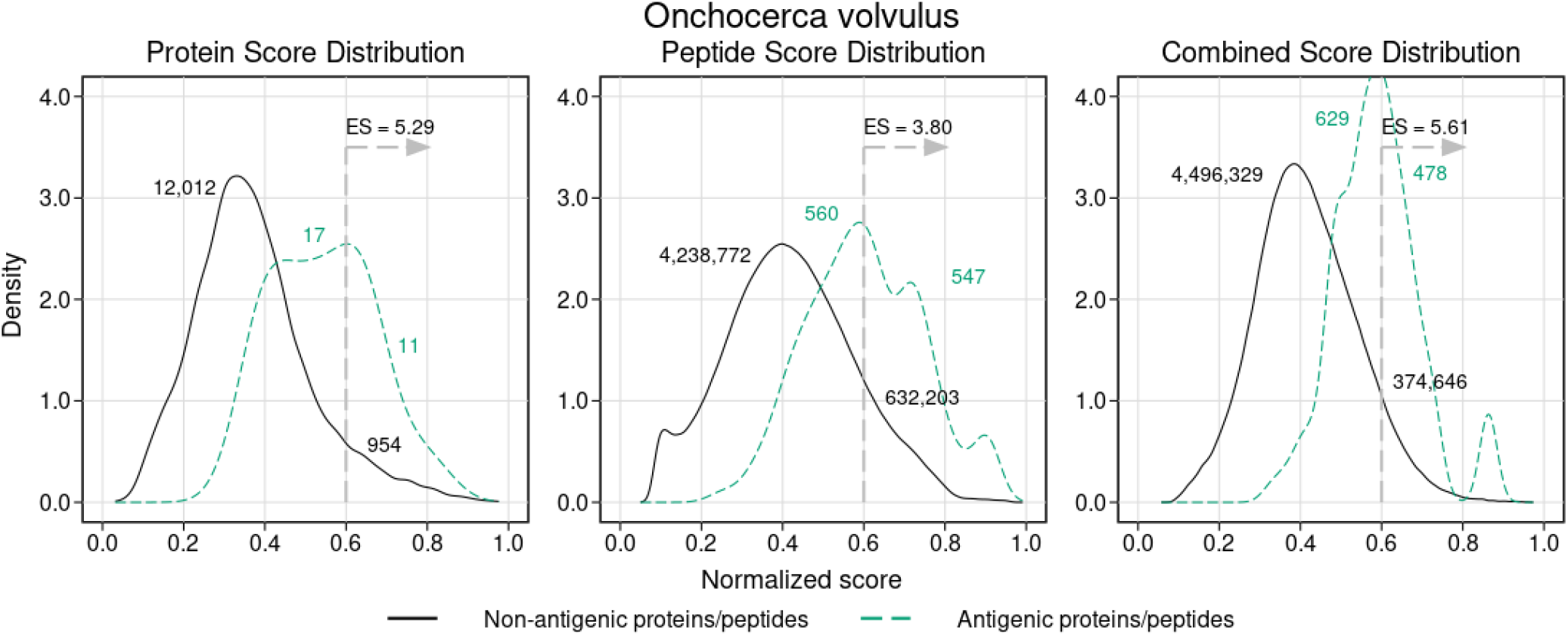
Density analysis for the antigenicity scores of *Onchocerca volvulus*. Plots were obtained by analyzing the proteome of *O. volvulus* with the final generic models, and then distinguishing between antigens and non-antigens. The figure shows the enrichment score obtained by keeping only the proteins and peptides with a score greater than 0.6, as well as the amount of antigens and non-antigens that would be inside or outside that subset. The plots correspond to the case where a protein was tagged as antigenic if it had at least 3 ‘immunoreactive’ peptides (see Results).

For peptides, APRANK obtained an enrichment score of 3.29 – 3.80, also showing an additive effect when combined with the protein score, suggesting that these are effective in predicting antigenicity for *O. volvulus*. Similar to before, we tried being more strict and only considering antigenic peptides in proteins with at least 2 or 3 ‘immunoreactive’ peptides; however this did not seem to affect the predictive performance as much as for whole proteins.

## 4 DISCUSSION

We present APRANK, a novel method to prioritize and predict the best antigen candidates in a complete pathogen proteome. APRANK relies on a number of protein features that can be calculated for any protein sequence which are then integrated in a pan-species model. Our benchmarks show that by integrating multiple predictors, pooling antigen data from multiple species across a wide phylogenetic selection, and balancing training datasets, APRANK matches or outperforms a state-of-the-art predictor such as BepiPred 1.0 in most scenarios.

We have tested this integrative method using non-parametric ROC-curves and made an unbiased validation using and independent data set (*O. volvulus*) containing recent proteome-wide antigenicity data. In summary, we found APRANK to be successful in predicting antigenicity for all pathogen species tested, hence providing a new and improved method to obtain antigen-enriched protein and peptide subsets for a number of downstream applications.

### 4.1 Conclusions: looking forward

While we are satisfied by APRANK’s performance, there are still ways to further improve it. The main issue we had when training our models is the current lack or sparsity of validated epitope and antigen information. Particularly, well validated non-antigenic sets are currently hard to find in the literature, forcing us to count as non-antigenic all proteins and peptides that do not currently have experimental evidence of antigenicity or were not tagged as antigenic in databases (which we know is hardly true).

We also observed that the performance of APRANK was not considerably affected by removing some individual features. This might indicate that, as we observed previously (Carmona et al., 2012), each individual predictor contributes only slightly to the overall performance. Another alternative explanation is that there might be redundancy between some of the predictors. For example the features being used for training of BepiPred 1.0 HMMs (propensity scales for secondary structure preference and hydrophilicity of amino acid residues (Larsen et al., 2006)) may overlap others used internally by some of the predictors in APRANK. Future versions of APRANK will review these overlaps, analyzing the pros and cons of adding novel predictors or removing existing ones.

Regarding the computing performance of APRANK, the majority of the time is dedicated to run the predictors used internally, most of which run in a reasonable time in a commodity server. However, there are a few bottlenecks (most notably predictions by NetSurfP). This should be improved in a future version in order to offer APRANK e.g. as a web-service. Future work will also explore the possibility to extend APRANK to also use data from other experimental (non-computable) sources, such as evidence of expression derived from proteomic or transcriptomic experiments.

### 4.2 Equations

See Supplementary Materials.

## CONTRIBUTION TO THE FIELD

The ability to predict which pathogen molecules elicit an immune response and are the target of antibodies during an infection is key for many diagnostic and clinical applications. Over time a number of predictors have been developed that seek to identify likely antigenic proteins and the portion of their structures that are recognized by antibodies (their epitopes). However this is a complex task which needs to be improved. Here we extend previous work and provide a new generalized method that succeeds in computing and extracting additional information from protein sequences, and use this information to train a model that can be used to prioritize candidate antigenic proteins from complete proteomes. Our integrative method –called APRANK– matches or outperforms existing predictors at the task of reducing the number of candidates down to a manageable and actionable number of likely antigenic proteins and epitopes. This is important for a number of downstream experimental assays. Using the described method and available software code, a complete pathogen proteome can be reduced to an enriched set of antigenic candidates for further evaluation.

## AUTHOR CONTRIBUTIONS

Conceptualization: ADR SJC FA; Data curation: ADR MB DR; Formal analysis: ADR MN FA; Funding acquisition: FA; Investigation: ADR; Methodology: ADR MB DR MN; Project administration: FA; Resources: FA; Software: ADR MB DR; Supervision: SJC FA; Validation: ADR; Visualization: ADR; Writing – original draft: ADR FA; Writing – review & editing: ADR MN FA.

## ACKNOWLEDGMENTS

We would like to thank Carlos Buscaglia (IIB-UNSAM-CONICET) for helpful discussions and critical review of the manuscript.

## SUPPLEMENTARY DATA

Supplementary Material should be uploaded separately on submission, if there are Supplementary Figures, please include the caption in the same file as the figure. LaTeX Supplementary Material templates can be found in the Frontiers LaTeX folder.

## DATA AVAILABILITY STATEMENT

The datasets analyzed for this study and the software used are available in this GitHub Repository: https://github.com/trypanosomatics/aprank. Trained models were deposited in Dryad under DOI:10.5061/dryad.zcrjdfnb1 (https://doi.org/10.5061/dryad.zcrjdfnb1).

## Supplementary Material

### 1 SUPPLEMENTARY DATA

Antigenic Proteins and Peptides used in this study were submitted separately as Supplementary Material. The corresponding file is an Excel spreadsheet containing complete listing of antigenic sources (proteins, peptides), their Uniprot and/or RefSeq identifiers and the corresponding mapping to our input sources (complete proteomes). File: ricci-aprank-supplementary-data.xlsx.

### 2 SUPPLEMENTARY TABLES AND FIGURES

#### 2.1 Figures

**Figure S1.**
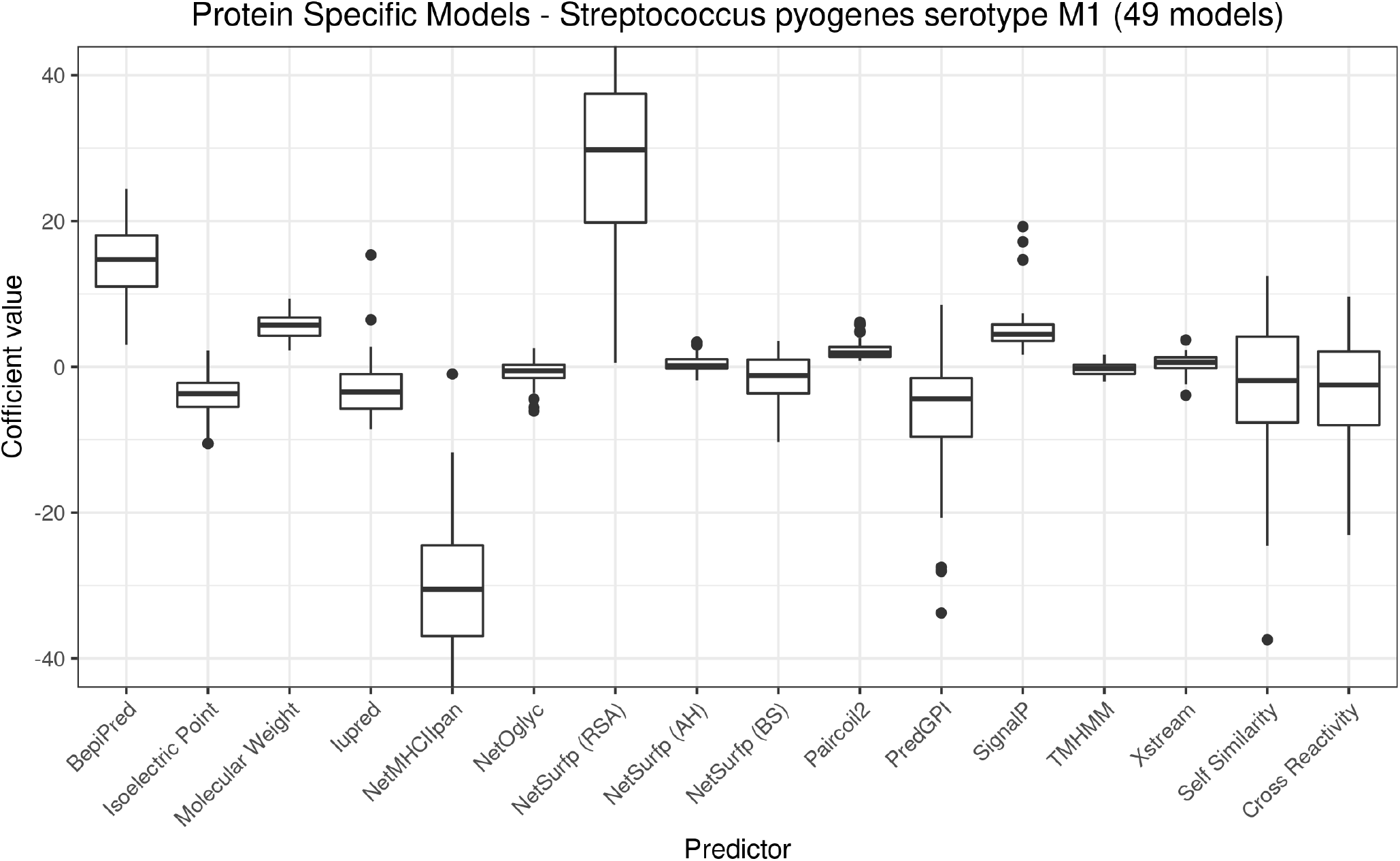
Coefficient values for the species-specific models for *Streptococcus pyogenes serotype M1*. Plots were obtained by recording the coefficient of each predictor in the binomial logistic regression models. These protein models correspond to the different species-specific models created when re-sampling training and test sets. One of the 50 models didn’t converge before reaching the maximum iteration limit when training, and so wasn’t considered.

**Figure S2.**
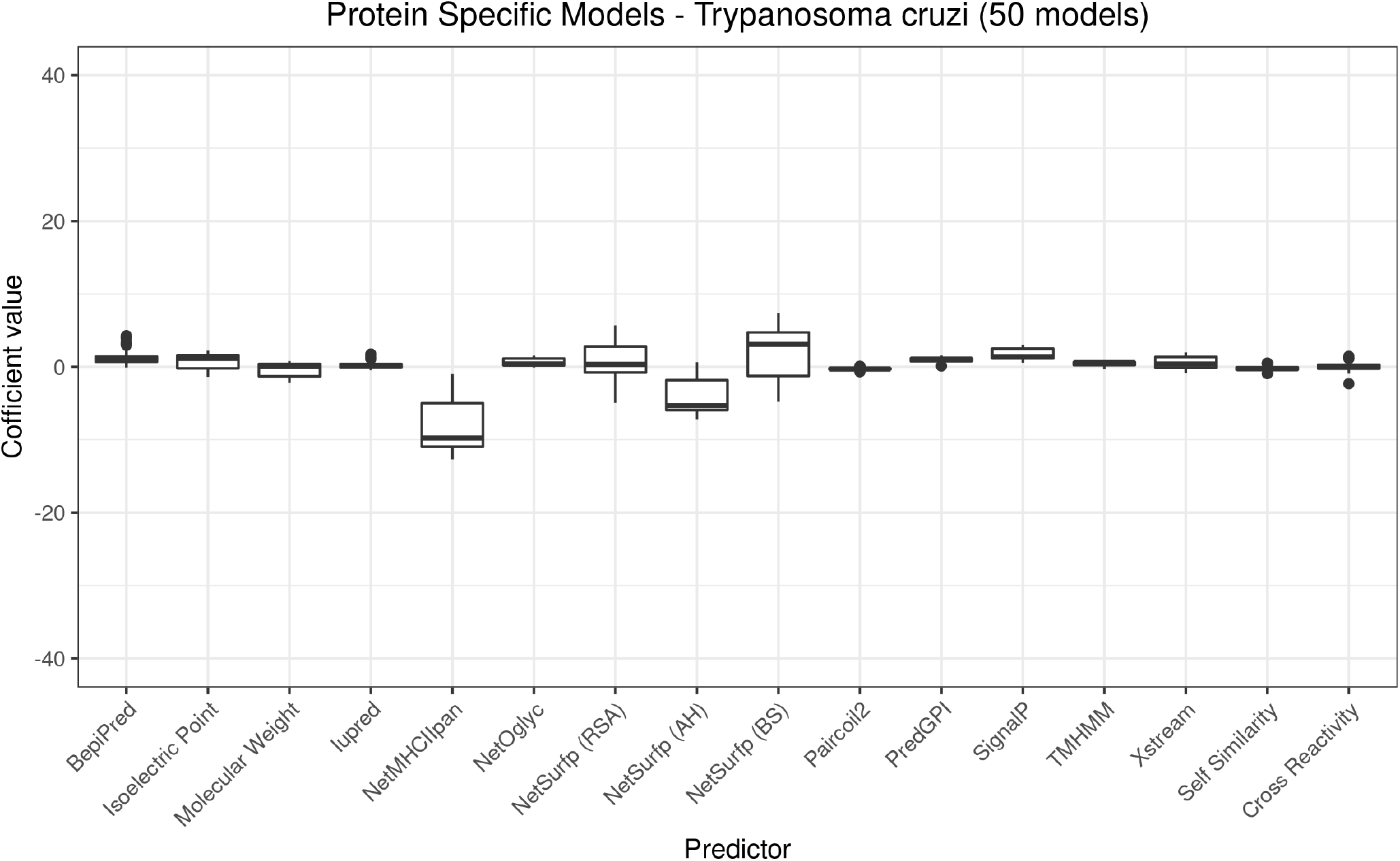
Coefficient values for the species-specific models for *Trypanosoma cruzi*. Plots were obtained by recording the coefficient of each predictor in the binomial logistic regression models. Different protein models correspond to the species-specific models created in each iteration when re-sampling training and test sets. All 50 models converged before reaching the maximum iteration limit when training.

**Figure S3.**
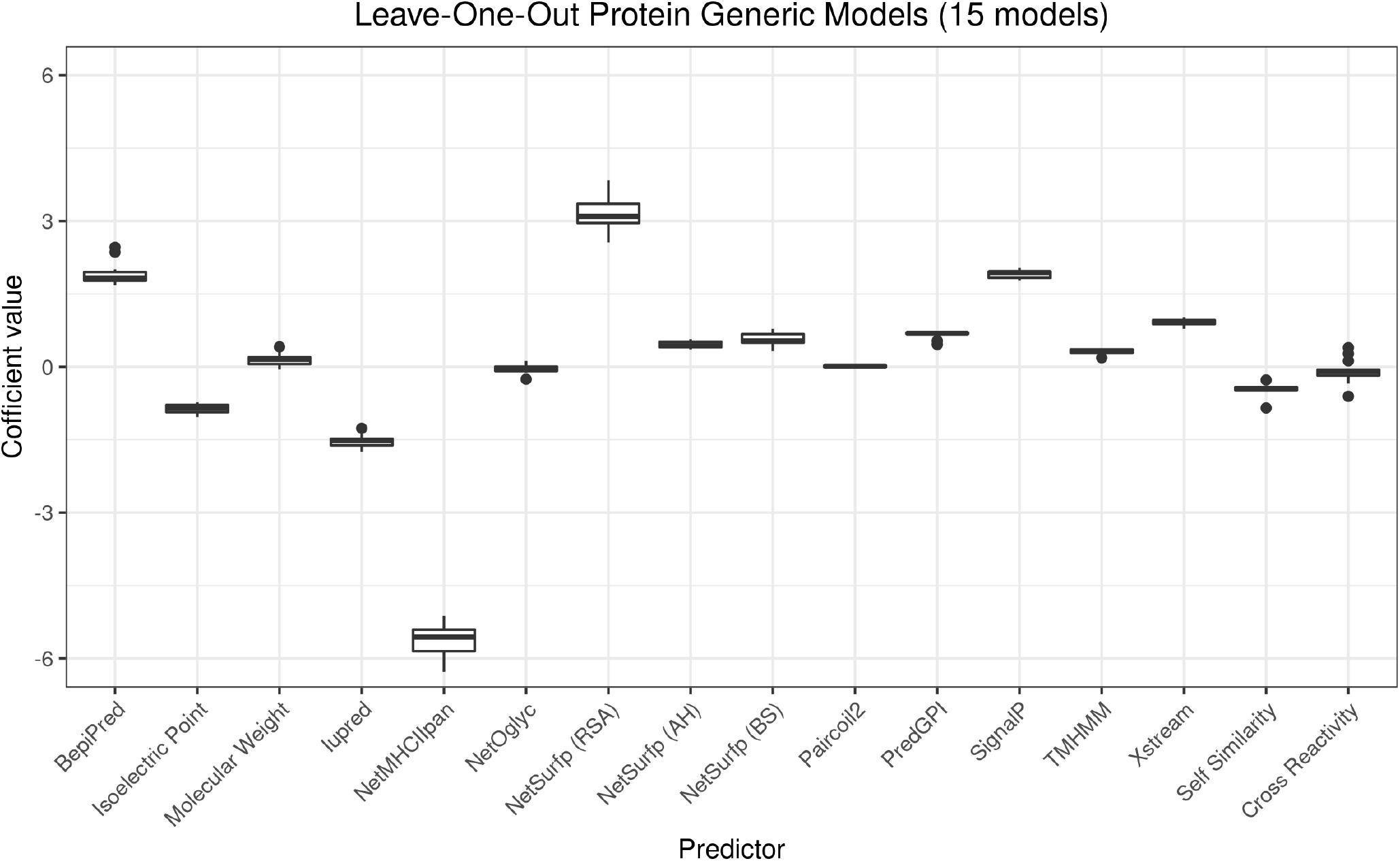
Coefficient values for the leave-one-out generic models. Plots were obtained by recording the coefficient of each predictor in the binomial logistic regression models. The different protein models correspond to each of the 15 leave-out-out generic models used to test APRANK. All 15 models converged before reaching the maximum iteration limit when training.

## 2.2 Tables

**Table S1.** Versions testing things of the software, packages and modules used to create our computational method.

| Software | Version |
| --- | --- |
| Ubuntu | 16.04 |
| R | 3.4.3 |
| ROSE (R package) | 0.0.3 |
| pROC (R package) | 1.12.1 |
| Perl | 5.22.1 |
| BioPerl (Perl module) | 1.007002 |

**Table S2.** Third-party software used to retrieve information about the proteins and peptides. The call being shown corresponds to those to use under Ubuntu 16.04. Words starting with $ symbolize variables to be replaced by their corresponding values.

| Predictor | Call | Data extracted |
| --- | --- | --- |
| BepiPred 1.0 | bepipred \$fasta_file -k >\$output_file | Score per amino acid |
| BLAST+ 2.2.31 | blastp -query \$query_file -db \$db_file<br>-outfmt 6 -out \$output_file<br>-max_target_seqs 2000 | Similarity between proteins (used to assign protein antigenicity) |
| EMBOSS 6.6.0.0 | pepstats -sequence \$sequence_file<br>-sprotein1 -aadata Eamino.dat -mwdata<br>Emolwt.dat -termini -nomono -auto<br>-outfile \$output_file | Isoelectric Point and Molecular Weight per protein |
| Iupred 1.0 | iupred \$sequence_file short >\$output_file | Score per amino acid |
| NetMHCIIpan 2.0 | netMHCIIpan -a \$allele -f \$sequence_file<br>-l \$peptide_length >\$output_file | %Rank per peptide per MHC II allele used |
| NetOglyc 3.1d | netOglyc \$sequence_file >\$output_file | Glycosilation presence per amino acid |
| NetSurfp 1.0 | NetSurfp \$sequence_file -a >\$output_file | Relative Surface Accessibility, Probability for Alpha-Helix and Probability for Beta-strand per amino acid |
| Paircoil2 | paircoil2 \$fasta_file \$output_file \$error_file | P-score per amino acid |
| PredGPI 1.4.3 | PredGPI.py \$filtered_fasta_file<br>>\$output_file | Presence and start of GPI per protein |
| SignalP 4.0 | signalp -f long -t \$organism_group<br>\$fasta_file \$output_file | Presence and start of signal peptide per protein and C and S score per amino acid |
| TMHMM 2.0c | tmhmm \$fasta_file >\$output_file | Presence of transmembrane helix per protein and amino acid participation in it per amino acid |
| Xstream 1.71 | java -jar \$Xstream_path/xstream.jar<br>\$sequence_file -d\$output_path/ | Start, end, period, copy number and consensus error per repeat per protein |

**Table S3.** Normalization methods used for each predictor in protein and peptide analysis. The formulas mentioned are shown in the supplementary materials.

| Predictor's output | Protein | Peptide |
| --- | --- | --- |
| BepiPred | Calculate the mean of the BepiPred score for the amino acids inside the protein and normalize it using fixedLinearNormalization with -1.5 and 1.5 as limits | Calculate the mean of the BepiPred score for the amino acids inside the peptide and normalize it using fixedLinearNormalization with -1.5 and 1.5 as limits |
| Isoelectric Point | Divide the isoelectric point value by 14 | - |
| Molecular Weight | Normalize the molecular weight value using sigmoidNormalization05 with a b of 30,000. | - |
| Iupred | Calculate the ratio of amino acids inside the protein with an score greater or equal than 0.5 | Calculate the ratio of amino acids inside the peptide with an score greater or equal than 0.5 |
| NetMHCIIpan | Calculate the mean of the ranks for the kmers inside the protein, divide it by 100, normalize it using fixedLinearNormalization with 0.05 and 0.5 as limits, and then subtract that number from 1 | Calculate the mean of the ranks for the kmers inside the peptide, divide it by 100, normalize it using fixedLinearNormalization with 0.05 and 0.5 as limits, and then subtract that number from 1 |
| NetOglyc | Calculate the ratio of glycosylated amino acids inside the protein, and normalize it using fixedLinearNormalization with 0 and 0.05 as limits | Check if the peptide has at least 1 glycosylated residue |
| NetSurfp (RSA) | Calculate the mean of the values for the amino acids inside the protein | Calculate the mean of the values for the amino acids inside the peptide |
| NetSurfp (Alpha Helix) | Calculate the mean of the values for the amino acids inside the protein | Calculate the mean of the values for the amino acids inside the peptide |
| NetSurfp (Beta Strand) | Calculate the mean of the values for the amino acids inside the protein | Calculate the mean of the values for the amino acids inside the peptide |
| Paircoil2 | Check if the protein has an amino acid sequence of a given length (50 by default) where all the amino acids has a score above a threshold (0.5 by default) | Calculate the ratio of amino acids inside the peptide with a score above a threshold (0.5 by default) |
| PredGPI | Check if the protein has a GPI | Check if the peptide is at least in part inside the GPI |
| SignalP | Check if the protein has a signal peptide | Check if the peptide is at least in part inside the signal peptide |
| TMHMM | Use the output as it is | Check if the peptide is at least in part inside a transmembrane helix |
| Xstream | Find the largest copy number in the protein and normalize it using sigmoidNormalization09 with a b of 5 | Assign to each amino acid the highest copy number it's involved in, calculate the mean of that value for the amino acids inside the peptide and normalize it using sigmoidNormalization09 with a b of 5 |
| Cross Reactivity | For each kmer in the protein, normalize the amount of times that kmer appears in the host proteome using sigmoidNormalization05 with a b of 1, then calculate the mean of these values | For each kmer in the peptide, normalize the amount of times that kmer appears in the host proteome using sigmoidNormalization05 with a b of 1, then, calculate the mean of these values |
| Self Similarity | For each kmer in the protein, normalize the amount of other times that kmer appears in the proteome using sigmoidNormalization05 with a b of 1, then calculate the mean of these values | For each kmer in the peptide, normalize the amount of other times that kmer appears in the proteome using sigmoidNormalization05 with a b of 1, then, calculate the mean of these values |

**Table S4.** Comparison between APRANK and a version of APRANK without the the predictor with highest solo AUC (BepiPred 1.0). The relative AUC gain shows the increase or decrease of the AUC obtained by APRANK relative to the version of APRANK without BepiPred. In bold we show differences greater than 5%. Due to the large number of peptides, each individual peptide AUC was calculated as the mean of 5 pseudo-random subsets of 50,000 peptides (see Methods).

| Species | Group | Proteins APRANK score |  |  | Peptides APRANK score |  |  |
| --- | --- | --- | --- | --- | --- | --- | --- |
|  |  | without BepiPred AUC | with BepiPred AUC | Relative AUC gain | without BepiPred AUC | with BepiPred AUC | Relative AUC gain |
| B. burgdorferi | Gram - | 0.777 | 0.786 | 1.18% | 0.726 | **0.768** | **5.78%** |
| B. melitensis | Gram - | 0.749 | 0.774 | 3.39% | - | - | - |
| C. burnetii | Gram - | 0.616 | 0.620 | 0.61% | - | - | - |
| E. coli | Gram - | 0.751 | 0.754 | 0.42% | 0.743 | 0.742 | -0.07% |
| F. tularensis | Gram - | 0.714 | 0.698 | -2.15% | - | - | - |
| L. interrogans | Gram - | 0.938 | 0.947 | 0.96% | 0.646 | **0.679** | **5.15%** |
| P. gingivalis | Gram - | 0.847 | 0.854 | 0.75% | 0.626 | **0.665** | **6.19%** |
| M. leprae | Gram + | 0.750 | 0.758 | 1.04% | 0.657 | **0.692** | **5.37%** |
| M. tuberculosis | Gram + | 0.697 | 0.702 | 0.66% | 0.586 | 0.586 | 0.00% |
| S. aureus | Gram + | 0.762 | 0.737 | -3.31% | 0.751 | 0.752 | 0.19% |
| S. pyogenes | Gram + | 0.983 | 0.983 | 0.04% | 0.826 | 0.838 | 1.47% |
| L. braziliensis | Eukaryote | 0.687 | 0.709 | 3.20% | 0.928 | 0.946 | 1.88% |
| P. falciparum | Eukaryote | 0.801 | 0.807 | 0.84% | 0.753 | 0.748 | -0.73% |
| T. gondii | Eukaryote | 0.835 | 0.837 | 0.27% | 0.585 | 0.583 | -0.47% |
| T. cruzi | Eukaryote | 0.869 | 0.867 | -0.29% | 0.833 | 0.843 | 1.26% |

**Table S5.** Comparison between APRANK and a ‘BLAST model’. The ‘BLAST model’ worked by assigning to each protein a score related to how similar they were to a recorded antigenic protein. For the two species that resulted in a better prediction when using the ‘BLAST model’, we also tested removing from the BLAST results (and so, from the ‘model’) the species that was the most similar to the one being analyzed. In bold we show differences greater than 5%.

| Species | Proteins |  |  |
| --- | --- | --- | --- |
|  | BLAST AUC | APRANK AUC | Relative AUC gain |
| B. burgdorferi | 0.502 | 0.786 | **56.60%** |
| B. melitensis | 0.637 | 0.774 | **21.49%** |
| C. burnetii | 0.579 | 0.620 | **7.14%** |
| E. coli | 0.677 | 0.754 | **11.44%** |
| F. tularensis | 0.629 | 0.698 | **10.92%** |
| L. interrogans | 0.499 | 0.947 | **89.86%** |
| P. gingivalis | 0.544 | 0.854 | **57.04%** |
| M. leprae | 0.893 | 0.758 | **-15.10%** |
| M. tuberculosis | 0.591 | 0.702 | **18.78%** |
| S. aureus | 0.622 | 0.737 | **18.56%** |
| S. pyogenes | 0.542 | 0.983 | **81.26%** |
| L. braziliensis | 0.951 | 0.709 | **-25.42%** |
| P. falciparum | 0.594 | 0.807 | **35.77%** |
| T. gondii | 0.443 | 0.837 | **88.98%** |
| T. cruzi | 0.501 | 0.867 | **72.95%** |
| M. Leprae (without M. tuberculosis in BLAST) | 0.650 | - | **16.63%** |
| L. braziliensis (without T. cruzi in BLAST) | 0.744 | - | -4.74% |

## 3 FORMULAS

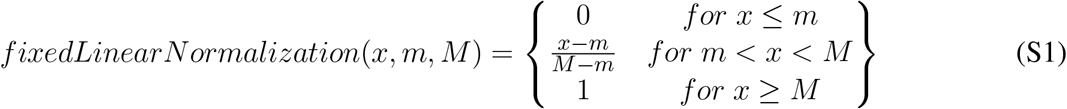

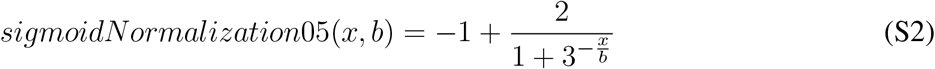

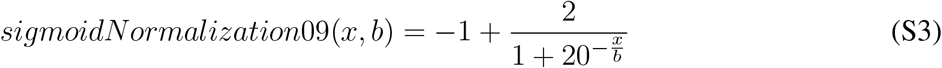

